# Competence to epithelialise coincides with competence to differentiate in pluripotent cells

**DOI:** 10.1101/809467

**Authors:** Chia-Yi Lin, Tulin Tatar, Guillaume Blin, Mattias Malaguti, Rosa Portero Migueles, Hongyu Shao, Naiming Chen, Ian Chambers, Sally Lowell

**Affiliations:** MRC Centre for Regenerative Medicine, Institute for Stem Cell Research, School of Biological Sciences, University of Edinburgh, 5 Little France Drive, Edinburgh EH16 4UU

## Abstract

Pluripotent cells reorganise themselves into an epithelium before they initiate differentiation, but it is not clear how these two events are mechanistically linked. Here we use quantitative imaging approaches to measure cellular rearrangements that accompany exit from naive pluripotency. We show that competence to epithelialise, like competence to differentiate, is a regulated process. The pro-differentiation transcription factor Tcf15 prospectively identifies cells that are competent to epithelialise. We identify early upregulation of the laminin receptor integrin alpha3 prior to differentiation and show that Tcf15 helps to regulate this change. Finally, we show that Tcf15 identifies and is required for efficient differentiation of a primed subpopulation of pluripotent cells. We conclude that competence to epithelialise is actively regulated and linked to differentiation-competence through the transcription factor Tcf15.

## Introduction

There is an increasing appreciation that morphological changes are not simply a consequence of differentiation but rather the two processes are often reciprocally interlinked (Gilmour *et al*, 2017; Chan *et al*, 2017). For example, in all amniotes pluripotent cells organise themselves into an epithelium prior to gastrulation (Nowotschin & Hadjantonakis, 2010; Bedzhov & Zernicka-Goetz, 2014; Sheng, 2015), and it has been suggested that this morphological change may be a prerequisite for subsequent lineage commitment (Ranga *et al*, 2014). At the same time, major changes in gene expression set up a new transcriptional state that makes pluripotent cells competent to differentiate (Nichols & Smith, 2009; Buecker *et al*, 2014; Boroviak *et al*, 2015). It is not understood whether the process of epithelialisation is mechanistically linked to differentiation-competence.

Pluripotent cells polarise before they commit to a primed state (Shahbazi *et al*, 2017). Polarisation is initiated by integrin-mediated adhesion to the laminin matrix secreted by the emerging extraembryonic endoderm (Bedzhov & Zernicka-Goetz, 2014; Li *et al*, 2016) and subsequently stabilised by PTEN (Meng *et al*, 2017) and Integrin-linked-kinase (Sakai, 2003). However, several open questions remain. Is the initiation of epithelialisation a spontaneous response to matrix or a regulated process? If epithelialisation is a regulated process, is it coordinated with differentiation or are the two processes regulated independently? What are the transcription factors and adhesion molecules that influence competence to epithelialise and help coordinate it with differentiation?

Tcf15 is a bHLH transcription factor that regulates epithelialisation of the somites (Burgess *et al*, 1996). It is also expressed in a subset of the Nanog-low subpopulation of mouse embryonic stem cells and within the inner cell mass of pre-implantation embryos (Davies *et al*, 2013). Tcf15 is activated transcriptionally by FGF and Wnt signalling (Linker *et al*, 2005; Davies *et al*, 2013). Its activity is suppressed post-translationally by BMP signalling: BMP upregulates Id1, which sequesters the E-proteins that form essential heterodimerisation partners with Tcf15 (Wilson-Rawls *et al*, 2004). A constitutively activated (BMP-resistant) form of Tcf15 has pro-differentiation activity in pluripotent cells (Davies *et al*, 2013), but the mechanism by which Tcf15 promotes differentiation has not been reported. Tcf15 null embryos survive through gastrulation (Burgess *et al*, 1996) but it is not known whether Tcf15 contributes to morphogenesis or differentiation during exit from naive pluripotency.

Here, we use quantitative image analysis to measure emergence of epithelial organisation during exit from naive pluripotency in vivo and in vitro. We show that although naive pluripotent cells can polarise they are not fully competent to epithelialise in response to matrix, whereas primed pluripotent cells can efficiently form an epithelium. Given that Tcf15 is expressed as cells downregulate Nanog (Davies *et al*, 2013) and that Tcf15 is required for epithelialisation of the somites (Burgess *et al*, 1996) we speculated that it may regulate epithelialisation of pluripotent cells. We use Tcf15 reporters and gain/loss of function approaches to show that Tcf15 identifies cells that are primed to epithelialise and that Tcf15 regulates the timely emergence the alpha subunit of the laminin receptor integrin α3β1. We also find that Tcf15 is required for efficient differentiation of a primed subset of pluripotent cells.

We conclude that, competence to epithelialise is an actively regulated process that is initiated in parallel with, rather than as a consequence of, exit from naive pluripotency. These two parallel processes are coordinated in part by the transcription factor Tcf15, analogous with the role of Tcf15 to coordinate epithelialisation and differentiation in the somites (Burgess *et al*, 1996).

## Results

### Pluripotent cells co-align into an epithelium as they become prepare to differentiate

We developed quantitative imaging tools for measuring the emergence of epithelial organisation based on segmenting individual nuclei and centrosomes and measuring nucleus centrosome vectors relative to the cavity of the embryo (Fig 1A) (Burute *et al*, 2017) https://pickcellslab.frama.io/docs/use/features/segmentation/nc_assignments/. We used this approach to assess the extent to which pluripotent cells are organised into an epithelium before and after implantation of the embryo. We confirm that pluripotent cells in the blastocyst have no apparent epithelial organisation. Nucleus-centrosome vectors measured relative to the cavity show no trend towards any particular direction: the number of cells pointing towards the cavity (angle <90°C) is similar to the number of cells pointing away from the cavity (angle >90°C) (Fig 1 B-C). In contrast, in the postimplantation epiblast almost all pluripotent cells point towards the cavity (angles <90°C) (Fig 1 B,D). These measurements confirm that pluripotent cells tend to co-align into an epithelial organisation during peri-implantation development, as expected (Nowotschin & Hadjantonakis, 2010).

**Figure 1:**
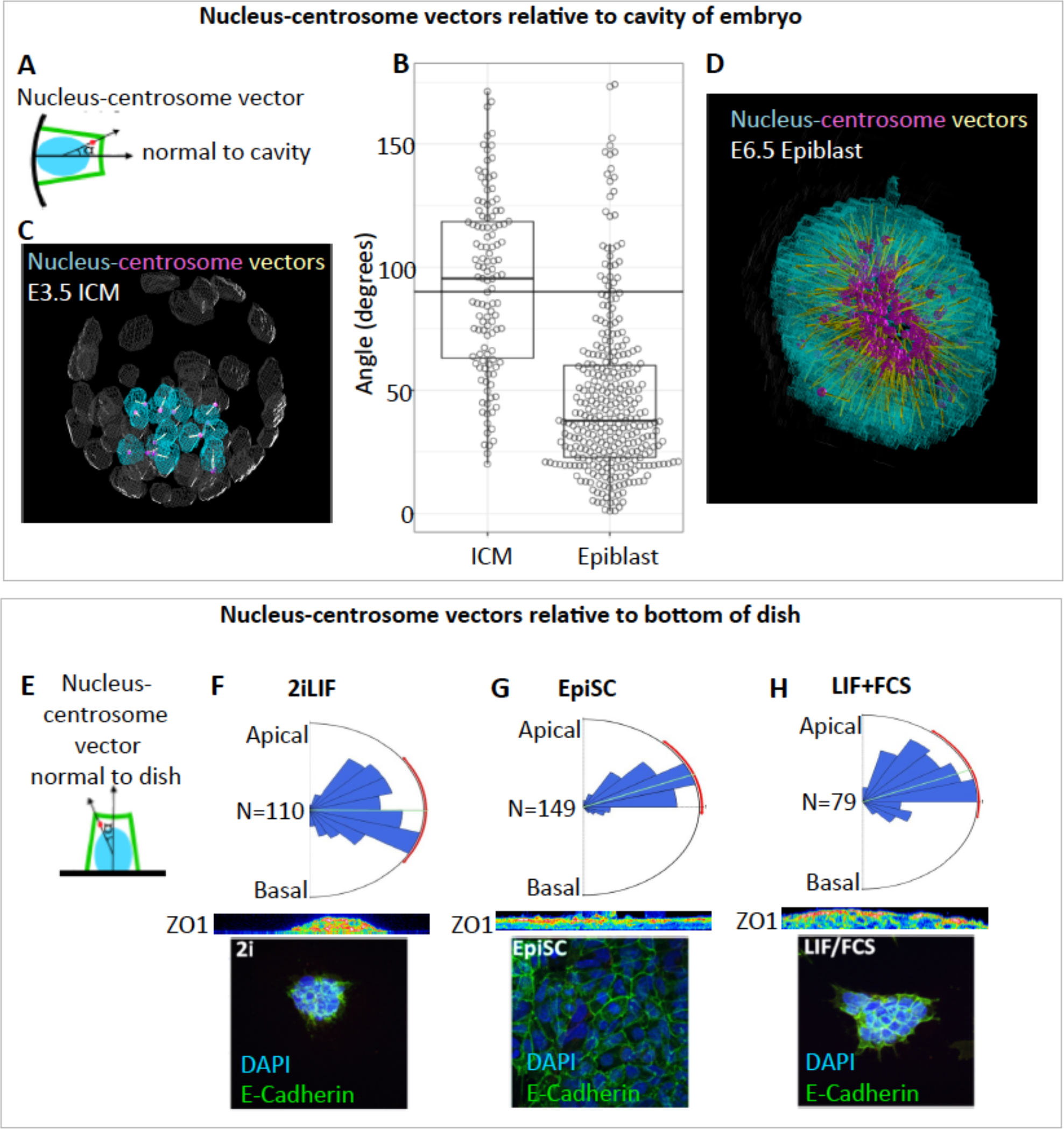
Pluripotent cells co-align into an epithelium as they prepare to differentiate. A-D: Cell polarity analysis in vivo. A: Schematic indicating the measurement of an angle between the nucleus to centrosome vector of a cell relative to the cavity of the embryo. B: Beeswarm-boxplot showing the distribution of angles between the nucleus and centrosome vector of the cell and its vector normal to the embryo cavity. Angles are shown for ICM cells in E3.5 blastocysts or for cells of the epiblast in E6.5 embryos. Angles are shown for ICM cells in E3.5 blastocysts (10 blastocysts for a total of 130 ICM cells) or for cells of the epiblast in one E6.5 embryo (354 cells). C-D: Representative 3D rendering of the processed images are given on each side of the plot. A shows an E3.5 blastocyst with and B shows an E6.5 embryo (transverse view). The nuclei of pluripotent cells are colored in cyan, extra-embryonic cell nuclei in grey, centrosomes in magenta and nucleus-centrosome vectors in yellow. E:H: Cell polarity analysis in pluripotent stem cell cultures. E: Schematic indicating the measurement of an angle between the nucleus to centrosome vector of a cell relative to the bottom of the dish F-G Rose diagrams show the distribution of angles between the nucleus to centrosome vector and its vector normal to the bottom of the dish. The green line in rose diagrams indicate the average angle, and the red arc represents the standard deviation of the angle. N indicates the number of cells included in the analysis. A representative confocal image is given for each culture condition stained for Zo1 to indicate AP polarity and E-Cadherin to demarcate cell outlines. ICM: inner cell mass

### Competence to epithelialise in response to matrix is acquired as pluripotent cells prepare to differentiate

In the embryo, laminin matrix is laid down by the emerging extraembryonic endoderm from late postimplantation development (Li *et al*, 2003). This raises the question of whether the formation of the pluripotent epithelium is a spontaneous response to the appearance of this extracellular matrix or whether it requires actively regulated intrinsic changes.

Formation of an epithelium involves multiple steps including acquisition of polarity, an increase in cell-matrix adhesion, and formation of a lumen, each of which could be independently regulated. It has previously been reported that naive pluripotent cells are able to polarise but unable to form a lumen (Shahbazi *et al*, 2017) and that polarisation occurs prior to neural differentiation (Ranga *et al*, 2014). In keeping with these reports, we confirm that naive cells do not efficiently form stable 3D cysts when cultured in matrigel but become competent to do so after 48h of neural differentiation in N2B27 (Supp Fig S1).

In order to analyse epithelialisation independently of luminogenesis, we turned to a 2D culture system. We analysed nucleus-centrosome vectors relative to the bottom of the culture dish (Fig 1E). We first analysed pluripotent cells grown in 2iLIF (Fig1F), which are equivalent to pluripotent cells of the preimplantation embryo (Boroviak *et al*, 2015) and compared these with epiblast stem cells (Fig 1G), which are equivalent to pluripotent cells of the postimplantation epiblast (Tesar *et al*, 2007; Brons *et al*, 2007). Each of these two cell types adopted an organisation similar to their in-vivo equivalents: 2iLIF cells fail to co-align while EpiSC co-align into an epithelial organisation with nucleus-centrosome vectors pointing up away from the surface of the dish (Fig 1F, G). These measurements confirm that cells co-align during the transition from naive to primed states in culture, as they do in vivo.

We then analysed cells cultured in LIF+FCS, where cells can exist in a mixture of naive and primed states within the same environmental conditions. Analysis of nucleus-centrosome vectors confirmed that, unlike naive cells, a majority of cells are aligned into an epithelium with centrosome pointing away from the bottom of the dish, but unlike primed cells a significant minority do not adopt this epithelial organisation (Fig1H): LIF+FCS). We conclude that pluripotent cells do not become fully competent to epithelialise until they reach a primed state.

### Tcf15 identifies cells that are competent to epithelialise

Tcf15 is a transcription factor that is expressed in the somites where it regulates epithelialisation (Burgess *et al*, 1996). Tcf15 is also expressed in the peri-implantation embryo and in a subset of pluripotent cells in LIF+FCS culture ((Davies *et al*, 2013) and Fig 2A) We therefore asked whether Tcf15 might identify the subset of pluripotent cells in LIF+FCS that are competent to epithelialise.

**Figure 2:**
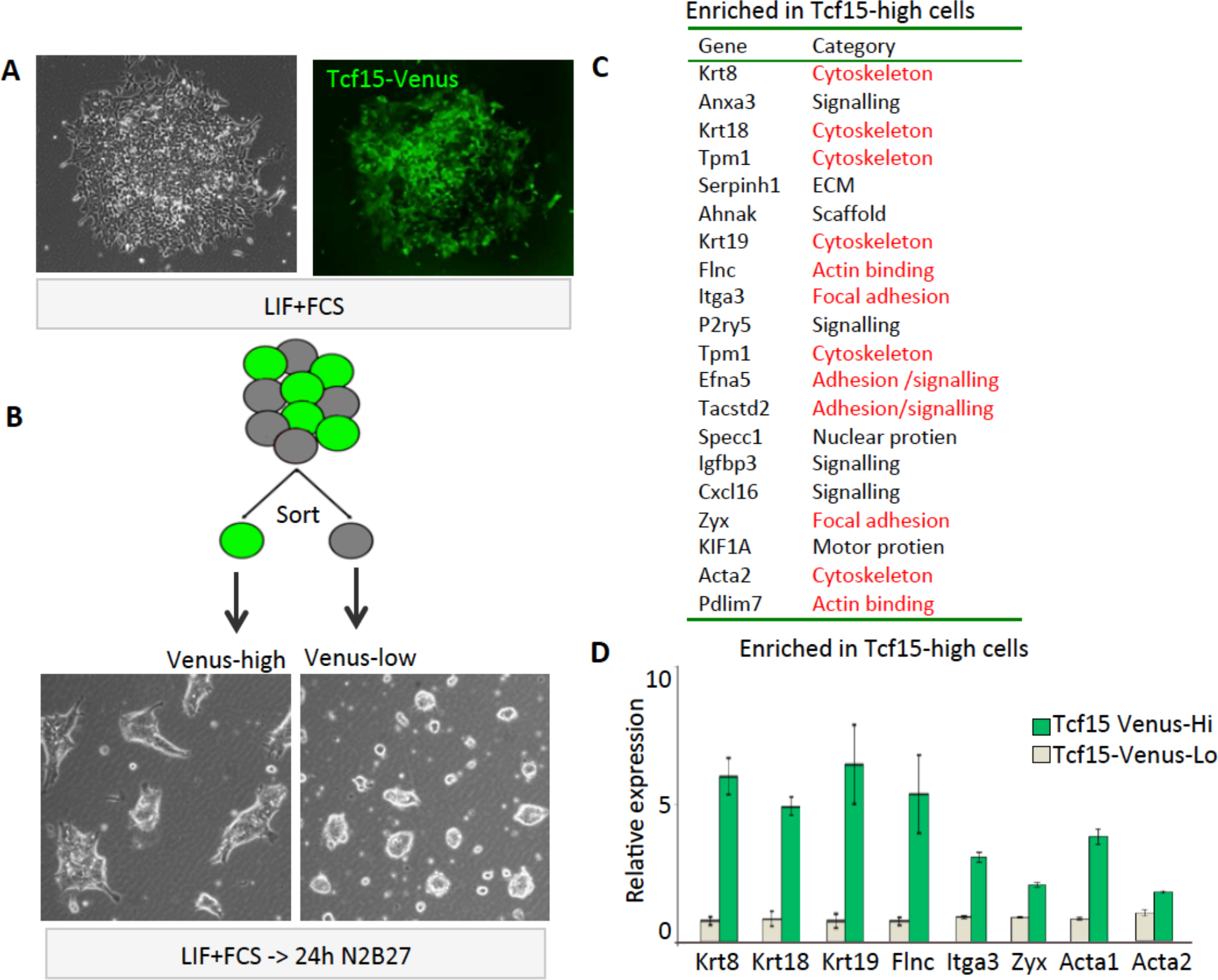
Tcf15 identifies cells that are competent to epithelialise. A: A typical colony of 216D1 Tcf15-Venus reporter ES cells expanded from a single cell in LIF+FCS for 6 days showing heterogeneous expression of Tcf15-Venus B: 216D1 Tcf15-Venus ES cell were cultured in LIF+FCS then sorted by FACS for Venus-high (top 30%) and Venus-low (bottom 30%) before plating on laminin for 24h in N2B27 media. Venus-high cells form flat epithelial colonies while Venus-low cells form domed colonies. C: 216D1 Tcf15-Venus ES cell were cultured in LIF+FCS then sorted by FACS for Venus-high (top 30%) and Venus-low (bottom 30%) before being subjected to microarray analysis: the top 20 differentially expressed genes between the two populations are listed here. Genes associated with adhesion and morphology are highlighted in red text. D: qPCR analysis of gene expression in FACS sorted Tcf15-Venus-high and Tcf15-Venus-low populations.

Using Tcf15-Venus reporter cell line 216D1 (Tanaka *et al*, 2008; Davies *et al*, 2013), we found that FACS-sorted Tcf15-Venus high cells form flatter more epithelial colonies than Tcf15-low cells when plated for 24h on laminin (Fig 2B). In order to explore the basis for this difference we performed transcriptome analysis on the sorted populations. We found that the top 20 genes enriched in Tcf15-Venus high cells included 12 genes encoding cell adhesion or cytoskeletal proteins (Fig 2C). Eight of these differences were confirmed by qPCR (Fig 2D). Of these, Integrin α3 (Itga3) is of particular interest because this forms part of the receptor for Laminin, which is the ECM to which pluripotent cells adhere during peri-implantation development (Li *et al*, 2003).

We conclude that Tcf15 identifies cells that form colonies with a distinct flattened morphology, and that genes encoding focal adhesion-associated molecules, including the laminin receptor Itga3 are enriched in Tcf15-high cells.

### Itga3 is upregulated as pluripotent cells prepare to differentiate

Having identified genes encoding adhesion and cytoskeletal proteins that are enriched in Tcf15-high cells, we asked if any of these genes are upregulated as naive cells progress towards a primed state.

We performed transcriptome analysis of cells at 12h after releasing cells from 2iLIF to EpiL conditions (FGF+Activin) (Fig 3A). Upregulated genes include read-outs of FGF signalling (Spry4, Dusp6) as expected, reflecting early responses to the change in extrinsic signals. Strikingly, Itga3 was the third most strongly upregulated gene at 12h (Fig 3A) in this genome-wide dataset.

**Figure 3:**
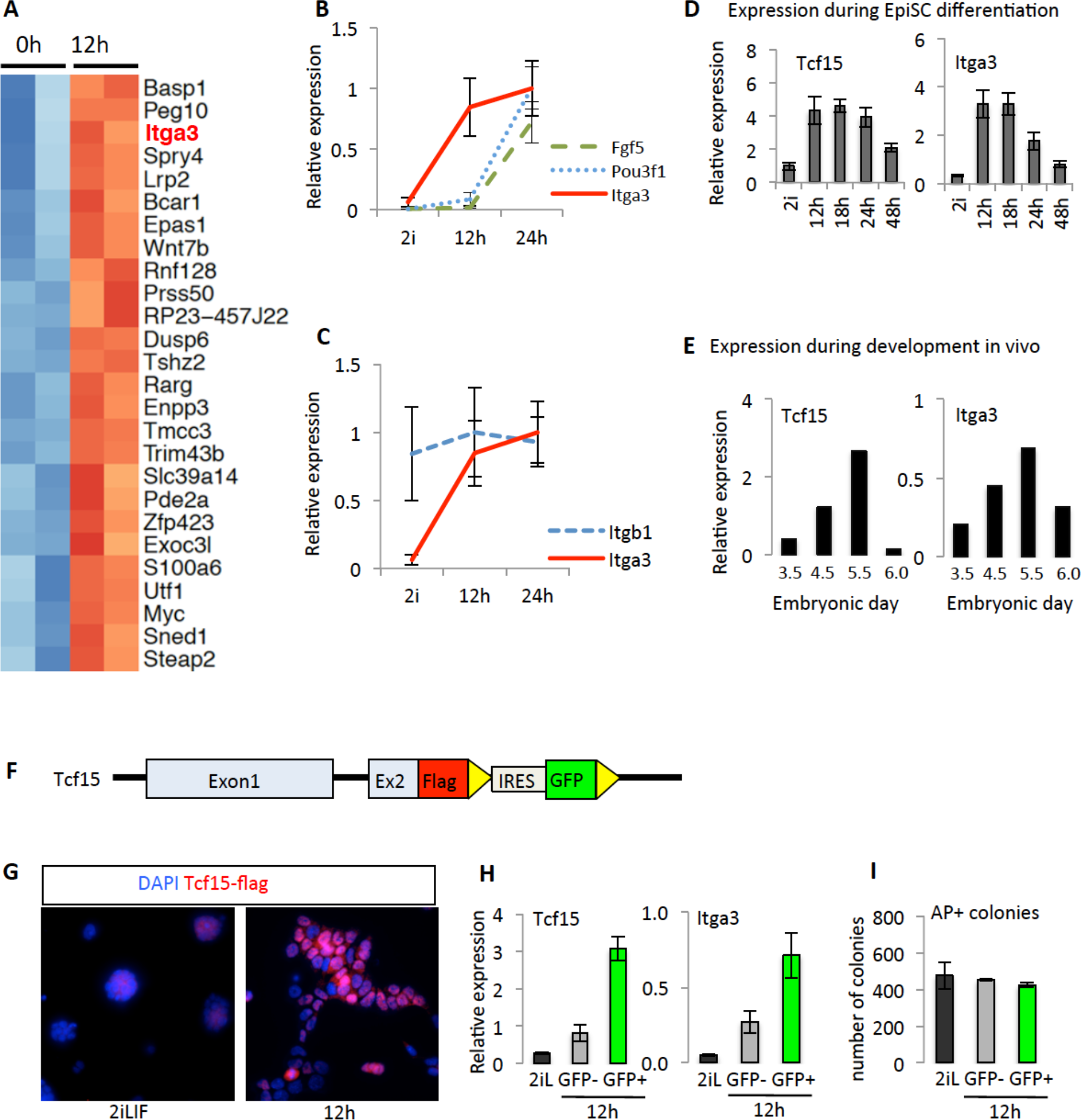
Itga3 is upregulated in Tcf15+ cells during the transition from naive to primed states. A: RNA-seq analysis on 0h (2iLIF) and 12h of EpiLC differentiation with E14Tg2a cells (in duplicates). Heat map represents the most strongly upregulated genes at 12h of EpiLC differentiation. B: qPCR analysis of *Fgf5, Pou3f1 and Itga3* mRNAs during EpiLC differentiation of E14Tg2a cells. Error bars represent the standard deviation of 3 independent experiments. C: qPCR analysis of *Itgb1 and Itga3* mRNAs during EpiLC differentiation of E14Tg2a cells (Error bars represent the standard deviation of 3 independent experiments). D: qPCR analysis of *Tcf15 and Itga3* mRNA levels during EpiSC differentiation of E14Tg2a cells. Error bars represent the standard deviation of 3 independent experiments. E: qPCR analysis of *Tcf15 and Itga3* mRNA levels at early stages of embryonic development. Embryos were collected from E3.5 (n=104, from outgrown culture for 4 days), E4.5 (n=108), E5.5 (n=126) and E6.0 (n=82). F: Graphical representation of *Tcf15* locus in Tcf15-(Flag)_3_ cells. Flag-IRES-GFP construct was inserted into the endogenous *Tcf15* locus at the 3’ end of the gene. Yellow triangles represent the loxP sites. G: Immunofluorescence analysis of Tcf15 protein in Tcf15-(Flag)_3_ cells during EpiLC differentiation. Tcf15 protein was detected with Flag antibody. H: Tcf15-(Flag)_3_ cells were differentiated under EpiLC conditions for 12h and then sorted based on GFP expression (which is a reporter for *Tcf15* transcription). *Tcf15* and *Itga3* transcript levels were detected by qPCR in sorted (GFP+ & GFP−) populations and starting 2iLIF population. I: Tcf15-(Flag)_3_ cells were differentiated under EpiLC conditions for 12h and then sorted based on GFP expression. Sorted cells were plated back into 2iLIF condition at clonal density and cultured for 6 days. The graph shows the colony numbers counted based on alkaline phosphatase staining. Error bars represent the standard deviation of 2 independent clones with 3 technical replicates each.

Upregulation of Itga3 occurs before cells have initiated expression of ‘primed’ markers Fgf5 or Pou3f1 (Fig 3B). Itga3 heterodimerises with Itgb1 to form the laminin receptor: in contrast to Itga3, transcripts for Itgb1 are readily detectable in naive cells and remain constant during the transition (Fig 3C).

We next confirmed that changes in expression of Itga3 follow a similar pattern to changes in expression of Tcf15 over the course of EpiSC differentiation (Fig 3D). It has previously been reported that Itga3 protein is detected in the epiblast of postimplantation embryos but not in the preimplantation blastocyst (Sutherland *et al*, 1993) We performed qPCR on embryos isolated before (E3.5) during (E4.5 - E5.5) and after (E6.0) implantation and confirmed that Itga3, similarly to Tcf15, is upregulated during peri-implantation development (Fig 3E).

We next asked whether the onset of Itga3 transcription correlates with the emergence of Tcf15 protein at the single cell level. While naive cells in 2iLIF are considered a homogenous population, differences emerge within 12h of placing cells under differentiation conditions (Kalkan *et al*, 2017). In order to examine the emergence of Tcf15 we engineered a cell line in which a flag epitope is fused to the endogenous Tcf15 protein while GFP reports Tcf15 transcriptional activity (Fig 3F) (Tatar et al in prep). We confirm that Tcf15 is barely detectable in naive cells but that Tcf15 protein (Flag staining) becomes detectable in a subset of cells within 12 hours of removing 2iLIF (Fig 3G). We sorted Tcf15-GFP high vs. low cells at 12h after removing 2iLIF and found that Itga3 transcripts are enriched within the Tcf15+ subpopulation within 12h of initiating exit from naive pluripotency (Fig 3H). There was no difference in the ability of the sorted subpopulations to form alkaline phosphatase positive colonies when plated at clonal density (Fig 3I), suggesting that cells have not yet irreversibly committed to a primed state at this 12h timepoint.

We conclude that the alpha subunit of the laminin receptor a3b1 integrin becomes upregulated within Tcf15+ cells prior to irreversible exit from naive pluripotency

### Tcf15 regulates changes in adhesion at the onset of differentiation

We next asked whether Tcf15 is responsible for controlling changes in adhesion at the onset of differentiation. We generated two independent Tcf15 null ES cell lines by deletion of the first Tcf15 exon, which encodes most of the coding region (Supp Fig 2 A-D).

Tcf15 null cells showed no gross defects when maintained in 2iLIF on laminin or in LIF+FCS on gelatin. They express normal levels of pluripotency markers and are able to differentiate into EpiLC or neural progenitors (Supp Fig 3A,B). They are also able to organise into epithelial cysts in matrigel with nucleus-centrosome vectors oriented appropriately towards the cavity, with only subtle differences from wild type cells (Fig 4A,B). We conclude that there are no major defects in the ability of Tcf15 null cells to polarise or to form a lumen, in keeping with the fact that Tcf15 null embryos are able to progress through development. However we noticed that Tcf15 null cell colonies adopted an unusual rounded morphology when plated on laminin (Fig 4C). In keeping with this phenotype, Tcf15 null cells cultured in LIF+FCS had lower expression of Itga3 (Fig 4D) and a reduced ability to upregulate Itga3 during the transition from naive to primed states (Fig 4E)

**Figure 4:**
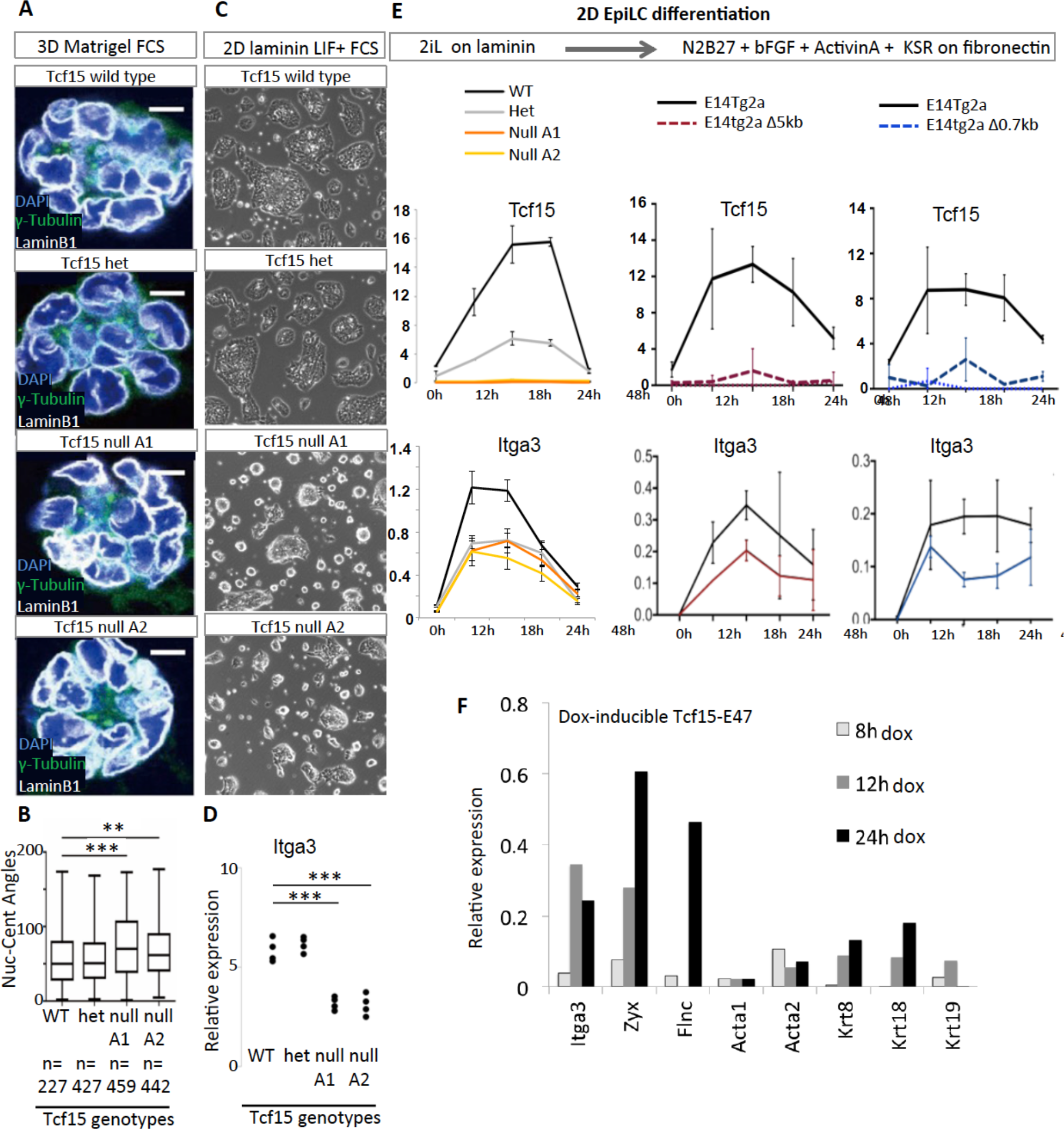
Tcf15 regulates changes in adhesion at the onset of differentiation. A: ES cells of stated genotypes were seeded at a density of 5000 cells/cm^2^ in GMEM+10% FCS in the absence of LIF and cultured for 48h before fixing and staining with DAPI and with antibodies against LaminB2 to mark the nuclear envelope and γ-tubulin to mark centrosomes. B: Nucleus-centrosome vectors were calculated for cell aggregates described in A. *** = P<0.01; ** = P<0.1. C: ES cells of stated genotypes were seeded onto laminin coated tissue culture plastic at a density of 105 cells/cm^2^ in GMEM+FCS and cultured for 24h before fixing. D: qPCR analysis of Itga3 mRNA in ES cells of stated genotypes cultured in LIF+FCS. E: qPCR analysis of cell lines of stated genotypes undergoing differentiation into EpiL cells. F: Microarray analysis of genes responding to induction of a Tcf15-E47. Cells engineered for dox-inducible expression of Tcf15-E47 (TTE15) were cultured in LIF+FCS and stimulated with 1ug/ml doxycycline for the stated times. Graph shows the average of duplicate microarray experiments. Error bars: standard deviation of 3 independent experiments.

In order to confirm this result, we sought alternative strategies to eliminate Tcf15 from pluripotent cells. We first used CrispR-Cas9 to delete 5kb of a candidate Tcf15 enhancer and showed that this was sufficient to completely eliminate Tcf15 expression (Fig 4E and Tatar et al in prep). We call these cells E14tg2a Δ5kb. We then generated another cell line, named E14tg2a Δ0.7kb in which 0.7kb of the Tcf15 enhancer was deleted. This also greatly reduced Tcf15 expression (Fig 4E and Tatar et al in prep). We used these cell lines to confirm that upregulation of Itga3 was significantly abrogated in the absence of Tcf15 (Fig4E).

These data indicate that Tcf15 contributes to rapid upregulation of the alpha subunit of the laminin receptor during the transition from naive to primed pluripotent states.

In order to confirm that Itga3 responds to activation of Tcf15 we used a cell line that is engineered for dox inducible expression of an activated form of Tcf15 (Davies *et al*, 2013). We found that Itga3 responds rapidly to Tcf15 activation: it is increased within 8h of dox treatment, which is the time that exogenous Tcf15 protein first becomes detectable (Davies *et al*, 2013).

We conclude that Tcf15 regulates changes in Itga3 expression at the onset of differentiation.

### Tcf15 identifies cells that are primed to differentiate

Our findings above (Fig 2) suggest that Tcf15 identifies a subset of cells that are enriched in Itga3 and competent to epithelialise. We have previously reported that activation of Tcf15 can prime cells to differentiate. We next asked whether the Tcf15-high subpopulation differentiates more efficiently than the Tcf15 low subpopulation.

We sorted Tcf15-Venus high and low cells and plated cells into basal N2B27 media to trigger the onset of differentiation. Tcf15-high cells differentiated more efficiently into neural cells compared with Tcf15-low cells (Fig5A-B). There was no difference between the two sorted subpopulations in their ability to form alkaline phosphatase positive colonies when plated at clonal density in LIF+FCS (Fig5C), indicating that Tcf15-high cells are not already differentiated or irreversibly committed to differentiation. We conclude that Tcf15 high cells differentiate more efficiently than Tcf15 low cells.

**Figure 5:**
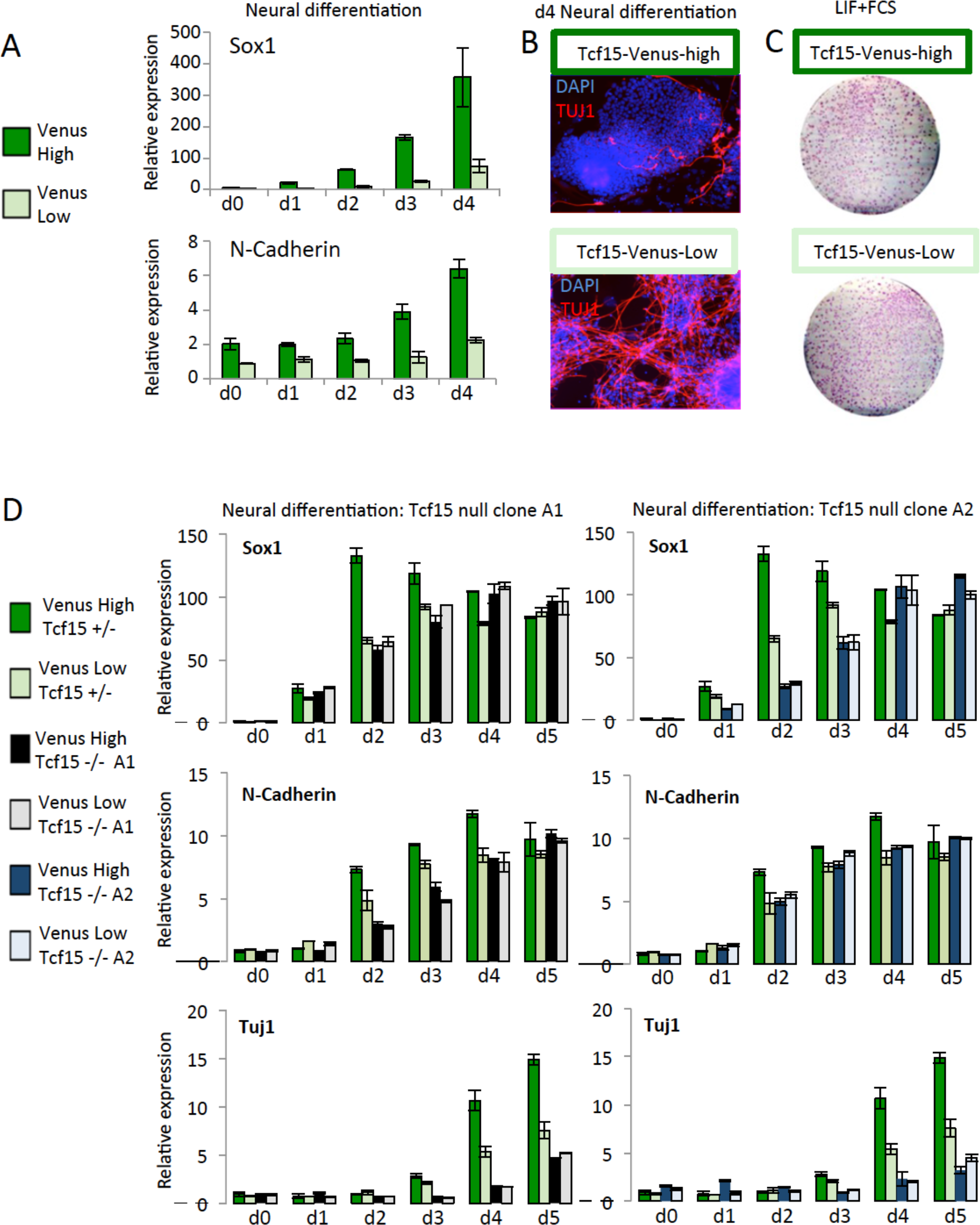
Tcf15 identifies cells that are primed to differentiate. A-B: 216D1 Tcf15-Venus reporter ES cells were sorted into Venus-high (top 30%) or Venus-low (bottom 30%) and plated into N2B27 to initiate neural differentiation. Samples were taken and analysed by qPCR each day for the next four days then analysed by qPCR for the stated markers (A) or stained for TUJ1 on day 4 (B). C 216D1 Tcf15-Venus reporter ES cells were sorted into Venus-high (top 30%) or Venus-low (bottom 30%) and plated at low density into LIF+FCS. Colonies were allowed to form over 6 days then the culture was fixed and analysed for alkaline phosphatase activity. D: Heterozygous or null Tcf15-VenusKI cells were sorted into Venus-high (top 30%) or Venus-low (bottom 30%) and plated into N2B27 to initiate neural differentiation. Samples were taken and analysed by qPCR each day for the next five days. Two different Tcf15 null clones were analysed: A1 and A2.

### Tcf15 is required for efficient differentiation of a primed subpopulation of pluripotent cells

Tcf15 is not absolutely required for development of mouse embryos through gastrulation nor for differentiation of pluripotent cells in bulk culture (Supp Fig 3). However, given that Tcf15 is able to prime pluripotent cells to differentiate (Davies *et al*, 2013) and that it becomes expressed in a subset of differentiation-primed cells upon exit from naive pluripotency (Fig2, Fig 5A-C) we speculated that Tcf15 may be specifically required for efficient differentiation of this primed subpopulation.

In order to address this we made use of heterozygous Tcf15 cells in which Venus replaces the first exon of Tcf15 (Tcf15VenusKI cells) (Supp Fig2). These cells have considerably reduced levels of Tcf15 protein (Supp Fig 2D), but were still able to detect a differentiation-advantage in the Tcf15-high subpopulation compared to the Tcf15-low subpopulation, as previously observed in our gene-conserving Tcf15 reporter cells (Fig 5D). When we deleted the second Tcf15 allele in order to eliminate Tcf15 expression completely, the differentiation-advantage was also eliminated: both the Venus-high and the Venus-low subpopulations differentiated at a similar rate to each other and no faster than Tcf15-low cells (Fig 5D). We conclude that the differentiation-advantage of Tcf15-high cells can be attributed directly to Tcf15 itself.

Overall, our data support a model in which the pro-differentiation transcription factor Tcf15 enables cells to respond efficiently to differentiation cues while at the same time allowing cells to rapidly upregulate Integrin a3 and epithelialise in response to laminin. Tcf15 is not absolutely required for either morphogenesis or differentiation, but contributes to the efficiency with which these two events are coordinated.

## Discussion

Amniotes organise their pluripotent cells into an epithelium as they prepare to gastrulate, suggesting that this morphological event may be an important prerequisite for the formation of the body plan (Sheng, 2015). Similarly, pluripotent cells in culture organise into epithelial sheets (in 2D culture) or epithelial cysts (in 3D culture) as they prepare to differentiate. Here we provide evidence that these morphological changes are not entirely a spontaneous response to extracellular matrix, but rather that they are actively regulated and linked to differentiation-competence through the transcription factor Tcf15.

The extrinsic events that drive epithelialisation are well described (Li *et al*, 2003), however the intrinsic changes that enable this process are less well understood. The epiblast forms through attachment of pluripotent cells to a laminin matrix that is secreted by the extraembryonic endoderm and the beta1 subunit of the laminin receptor, encoded by Itgb1, is required for epithelialisation of pluripotent cells(Bedzhov & Zernicka-Goetz, 2015). However Itgb1 is expressed in naive cells, which are not competent to fully epithelialise, suggesting that additional factors may be required. We find that Tcf15 upregulates the alpha subunit of the laminin receptor, Itga3. We show Itga3 is upregulated very early, prior to irreversible exit from naive pluripotency, in vitro and are also upregulated during peri-implantation development, in keeping with previous studies (Sutherland *et al*, 1993; Hayashi *et al*, 2011; Mohammed *et al*, 2017). However it is unlikely that these are the only changes driving epithelialisation: for example proteomic (Fröhlich *et al*, 2013) and transcriptomic (Hayashi *et al*, 2011) studies reveal that a broad set of actin-regulators and intermediate filaments are upregulated transition from naive to primed states in vitro. Morphological changes are therefore likely to be the result of a complex and coordinated change in the expression of adhesion and cytoskeletal regulators, some but not all of which are controlled by Tcf15.

The mechanism by which Tcf15 drives differentiation remains unclear. Tcf15 is able to rapidly downregulate Nanog (Davies *et al*, 2013), which might help to explain prodifferentiation effect, but the lack of a global differentiation defect in Tcf15 null cells (Supp Fig 3) or embryos (Burgess *et al*, 1996) argues that other factors must act redundantly with Tcf15 to trigger the loss of naive pluripotency transcription factors. Here we show that differentiation defect in Tcf15 null cells becomes apparent when Tcf15-high cells are assessed side by side with Tcf15-low cells. This suggests that Tcf15 has relatively subtle effects on the efficiency of differentiation, in keeping with a model in which Tcf15 drives changes in adhesion that then influence the robustness of the differentiation process.

This raises the question of whether Tcf15’s influence on adhesion and differentiation is regulated independently or whether adhesive changes might reinforce pro-differentiation signals. This is difficult to test because eliminating adhesion causes cell death in epiblast cells (Li *et al*, 2002; Migeotte *et al*, 2010), but it is tempting to speculate that differentiation competence may be enhanced by the ability of integrins to activate pro-differentiation signals (Humphries *et al*, 2018). This type of mechanism might help to coordinate differentiation with morphogenesis by providing a differentiation advantage to cells that have initiated epithelialisation.

There has been a great focus on the transcriptional and epigenetic events that prepare pluripotent cells to differentiate, but less appreciation of changes in adhesion and morphology that could influence differentiation (Malaguti *et al*, 2013; Livigni *et al*, 2013). While these changes in adhesion and morphology are unlikely to be dominant drivers of differentiation, they are likely to influence the efficiency with which cells can respond to differentiation cues and may be important for making development more robust. Here we provide one mechanistic link between differentiation and changes in adhesion that help drive morphological changes. Our increasing understanding of these links will help clarify the relationship between morphogenesis and differentiation in pluripotent cells

## Acknowledgements

We are grateful to the staff at the SCRM FACS facility, Tissue Culture facility, transgenic unit, and imaging facility for help and guidance. We thank Raphael Pantier, Dougie Colby, Darren Wisniewski Karolina Punovuori and Sam Wu and Daina Sadurska for help and advice and members of the lab for useful discussions. SL is funded by a Wellcome Trust Senior Fellowship WT103789AIA and GB was funded by a Wellcome Trust Sir Henry Wellcome Fellowship WT100133.

## Materials and methods

### Cell lines

Cell lines made in this study were generated from mouse ES cell line E14tg2a (Smith & Hooper, 1987).

Cell line ‘TTE15’ engineered for dox-inducible expression of a Tcf15-E47 forced dimer is described in (Davies *et al*, 2013). 1 μg/mL of doxycycline was used to induce expression of the transgene.

Cell line 216D1 is a Tcf15 reporter cell line which a *gtx-IRES-Venus* cassette was incorporated into the *Tcf15* locus downstream of the coding sequence (Tanaka *et al*, 2008). This cell line preserves both coding alleles.

Tcf15-VenusKI ES cells were generated by replacing the first exon of Tcf15 with the coding sequence for Venus. Tcf15-VenusKI +/− refers to cells after one round of targeting, i.e. heterozygous for the Tcf15 allele. Tcf15-VenusKI −/− refers to cells in which two rounds of targeting were performed in order to replace both alleles.

Sox1-GFP (Ying & Smith, 2003) were differentiated into EpiSC following the protocol described in (Guo *et al*, 2009).

All newly derived lines and clonal lines were confirmed to have a normal karyotype before being used in experiments.

### Cell culture

LIF+FCS culture of ES cells: Glasgow Modified Eagle’s Medium (GMEM), 10% foetal calf serum, 1 mM sodium pyruvate, 1× non-essential amino acids, 2 mM L-glutamine, 0.1 mM 2-mercapthoethanol and 100 units/mL LIF. LIF+FCS cells were grown on gelatin 2iLIF culture of ES cells: N2B27 supplemented with 3 μM CHIR99021, 1 μM PD0325901 and 100 units/ml LIF. 2iLIF cells were grown on laminin.

EpiSC were cultured in N2B27 supplemented with 12 ng/mL FGF2 and 20 ng/mL Activin A. EpiSC were grown on fibronectin. EpiLC differentiation was performed as previously described (Hayashi *et al*, 2011) and EpiSC differentiation was performed as previously described (Guo & Smith, 2010). Neural differentiation was performed in N2B27 media on gelatin as previously described (Pollard *et al*, 2006). N2B27 media consists of 50% Neurobasal, 50% DMEM/F12 (without L-glutamine), 0.1 mM 2-mercapthoethanol, 2 mM L-glutamine, 0.5% modified N2 supplement (made in house) and 1% B27 supplement.

Laminin was used at 5 μg/mL in PBS and dishes were coated for overnight at 4°C.

Fibronectin was used at 7.5 μg/mL in PBS and dishes were coated at room temperature for 10 minutes.

### Targeting construct for Tcf15

The targeting construct for Tcf15 contained a 2.3kb 5’ homology arm and a 2.4kb 3’ homology arm corresponding to the sequences flanking the first coding exon of Tcf15. Within the two homology arms is the coding sequence for Venus followed by an frt-flanked pgk-neo MC1-TK cassette selection cassette. This cassette was removed by transfection of pFlpO after isolation of successfully targeted clonal cell lines. A PGK-DTA cassette was placed upstream of the 5’ homology arm to enable negative selection of clones in which the targeting cassette had randomly integrated.

### Southern blotting

Genomic DNA was extracted from mouse ES cells using the DNeasy Blood and Tissue kit (Qiagen). Southern blotting was performed as previously described (Southern, 1975).

### pPCR primers

Acta1 tgaagcctcacttcctaccc, cgtcgcacatggtgtctagt

Acta2 ctctcttccagccatctttcat, tataggtggtttcgtggatgc

Fgf5 aaaacctggtgcaccctaga, catcacattcccgaattaagc

Flnc ggagtcctttccctgtccat, tcacacgcacacctttgg

Itga3 aggatatgtggcttggagtga, gaccacagcaccttggtgta

Itgb1 atgcaggttgcggtttgt, gggtaaaacaataccaccaagttt

Krt18 gaggcagagattgccaccta, agggcatcgttgagactga

Krt19 tgctggatgagctgactctg, cctcagggcagtaatttcctc

Krt8 agttcgcctccttcattgac, gctgcaacaggctccact

N-cadherin gccatcatcgctatccttct, ccgtttcatccataccacaaa

Otx2 ccacttcgggtatggacttg, gtcctctcccttcgctgttt

Pou3f1/Oct6 cgctgtatgctgtatttatcgg, atacacagatgcggctctc

Sox1 gtgacatctgcccccatc, gaggccagtctggtgtcag

TBP ggggagctgtgatgtgaagt, ccaggaaataattctggctca

Tcf15 gctccatctgcaccttctg, ggctacacccctcactttca

TUJ1 tcactgtgcctgaacttacc, ggaacatagccgtaaactgc

Zyx ccagggagaaagtgtgcagt, gttcttggtcatgtcgtcca

### Transcriptome analysis

For RNAseq analysis total RNA was isolated from cells using the Absolutely RNA Miniprep Kit (Agilent) and RNA quality was verified using an Agilent 2100 Bioanalyzer. Subsequent cDNA library preparation, sequencing and bioinformatics analysis, including differential gene expression analyses, were performed by the Edinburgh Genomics facility. Library preparation was performed using the TruSeq Stranded mRNA Library Prep Kit (Illumina) and libraries were sequenced on the Illumina HiSeq4000. Reads were mapped to the mouse genome GRCm38 from Ensembl and were aligned to the reference genome using STAR (version 2.5.2b). Differential gene analysis was carried out with edgeR (version 3.16.5). Gene ontology analysis was performed on genes upregulated at 24h using the STRING database (Szklarczyk *et al*, 2017).

For microarray analysis RNA was prepared using Absolutely RNA Purification Kits cRNA was synthesized using an Illumina TotalPrep RNA Amplification Kit. Labelled RNA was submitted to the WTCRF MRC Human Genetics Unit (University of Edinburgh) for further processing. cRNA quality was checked using an Agilent 2100 Bioanalyser and hybridization was performed on a MouseWG-6 v2 BeadChip (Illumina). Raw data was processed in R using the beadarray (Dunning *et al*, 2007) and limma (Smyth *et al*, 2005) packages from the Bioconductor suite (Gentleman *et al*, 2004). RNAseq data is available in GEO with accession number GSE132003 and microarray data is available in GEO with accession number GSE42539.

### Clonal self-renewal assays and alkaline phosphatase staining

Cells were plated 10-30 cells/cm^2^ in LIF+FCS and media was changed every other day. After 7 days, alkaline phosphatase staining was performed using the Leukocyte Alkaline Phosphatase Kit (Sigma).

### Fluorescence Activated Cell Sorting

Cells were dissociated into single cell suspensions in ice-cold PBS+10% FCS, in the presence of either 100ng/ml DAPI or 1μg/ml propidium iodide to stain dead cells. Cell sorting was performed on a BD FACSAria. Tcf15-Venus reporter cells were sorted for a Tcf15-Venus-high subpopulation by gating for the 30% of cells with the highest Venus fluorescence and for a Tcf15-Venus-low subpopulation by gating for the 30% of cells with the lowest Venus fluorescence. Cells were also gated for immunoreactivity to PECAM in order to exclude differentiated (PECAM-negative) cells.

### Embryo collection and chimera generation

Pre- and peri-implantation embryos were obtained by flushing uteri with a large-bore blunted needle in M2 medium (Sigma). Post-implantation embryos were dissected at 5.5 and 6.5 d.p.c. in M2 medium.

For generating chimeric mice, F1 female mice were superovulated (100 IU/ml PMSG, ProSpec, and 100 IU/ml HCG, Intervet, intraperitoneal injections 48h apart) and crossed with wild-type stud male mice. Pregnant mice were culled at 2.5 d.p.c. by cervical dislocation, ovaries with oviducts were dissected and collected in pre-warmed M2 medium. Oviducts were flushed using PBS and a 20-gauge needle attached to a 1ml syringe and filled with PB1. 2.5 d.p.c. embryos were collected and washed in PB1, the zona pellucida was removed using acidic Tyrode’s solution (Sigma), and transferred to a plate with incisions where one clump of 8-15 cells were added to each embryo. Embryos were then incubated for 24h at 37°C in 5% CO_2_.

Blastocysts were selected and collected to be transferred into the uterus of a pseudopregnant CD-1 female. Embryos were dissected at 10.5d.p.c. in M2 medium and observed for chimeric ES cell contribution.

Mice were housed and bred in the Animal Unit of the Centre for Regenerative Medicine, in accordance to the provisions of the Animals (Scientific Procedures) Act 1986 and the 2010/63/EU Directive.

### Western blotting

Cells were lysed in RIPA buffer + 1× PMSF (Alpha Diagnostics). 20μg protein lysates were run on 4-12% NuPage Bis-Tris Gel (Novex) and transferred onto Amersham Hybond ECL Nitrocellulose Membrane (GE Healthcare). Membranes were blocked in 5% Amersham ECL Prime Blocking Agent (GE Healthcare) + 0.1% Tween 20 (Sigma) in PBS. Membranes were incubated in primary antibody overnight at 4°C, washed 3 times in PBS + 0.1% Tween 20, incubated in HRP-conjugated secondary antibody for 1 hour at room temperature and washed 3 times in PBS + 0.1% Tween 20. The membrane was incubated in Amersham ECL Western Blotting Detection Reagent (GE Healthcare) or Amersham ECL Prime Western Blotting Detection Reagent (GE Healthcare), depending on the expected strength of signal. The membranes were used to expose Amersham Hyperfilm ECL (GE Healthcare), and films were developed using a Konica SRX-101A Medical Film Processor. The Tcf15 antibody used for Western blotting was obtained from Santa Cruz (sc-46438) and used at a dilution of 1:100.

### Immunofluorescence

Samples were fixed in 4% formaldehyde for 10 min at room temperature (cells and blastocysts) or for 30 min at 4°C (E6.5 embryos). The fixative was quenched with 50mM ammonium chloride (Sigma) dissolved in PBS/0.1%Triton for 5 min. The cells were then incubated for a minimum of 30 min (cell culture) or overnight (embryos) with blocking solution which consisted of 5% donkey serum (Sigma), 0.1% Triton X-100 (Sigma) and 0.03% Sodium Azide (Sigma) in PBS. Incubation with antibodies was performed for 1h (cell culture) or overnight (embryos) at room temperature in a humidified chamber. Antibodies were all diluted in blocking solution. Cells grown on glass coverslips were mounted in ProLong© Gold Antifade Mountant (Molecular Probes) 24h prior imaging. Embryos were further clarified using a Benzylalcohol/Benzylbenzoate (BABB) based method adapted from (Dodt et al., 2007). Briefly embryos were dehydrated in graded methanol series (25%, 50%, 75%, 90%, 100% twice) of 5 min each. Embryos were then transferred to a 1:1 solution of BABB/Methanol for 5 min before a final transfer into a pure solution of BABB prior imaging.

Nuclei were counter-stained with Dapi except when samples were used for image analysis. In this latter case, embryo samples were stained for LaminB1 to use as input for Nessys (Blin *et al*, 2019) and stained with propidium iodide to use as input for FARSIGHT (Al-Kofahi *et al*, 2010) (see image analysis section). Propidium iodide staining was performed as follows: After immunostaining, cells were incubated in 0,03M Na Citrate (pH7) and 0,3M NaCl (SSC buffer) for 5 min. The cells were then treated with 10μg/ml of RNAase A for 30 min in SSC at 37°C, rinsed once with SSC buffer and incubated with 1,5μM propidium iodide (in SSC). Finally, the cells were rinsed 3 times with SSC buffer and mounted in ProlongGold.

Antibodies used for immunofluorescence: LaminB1 1/1000 (Abcam ab16048, 1/1000), ZO-1 (Zymed 339100, 1/200), gamma-Tubulin (Abcam ab11316, 1/500), Ecad (Zymed 13-1900, 1/200), Ncad (BD 610920, 1/200), Flag (Sigma 1804 1/500), Tbra (R&D AF2085, 1/400), TUJ1 (Covance MMS-435P 1/500), Oct4 (Santa Cruz sc-5279 1/200). All secondary antibodies were Alex-Fluor conjugated (Invitrogen) and used at a dilution of 1/1000.

### Imaging

Pluripotent cells were imaged with a Leica SpE inverted confocal microscope using an ACS APO 63× objective with oil immersion and NA=1.3. Image voxel sizes: 0.085 × 0.085 × 0.42 μm.

Blastocysts and E6.5 embryos were imaged with a Leica TCS-Sp8 inverted confocal microscope using either an HC PL APO 40× oil objective with NA=1.3 for Blastocysts or a 25× water immersion objective with NA=0.95 for E6.5 embryos. Voxel sizes: 0,28 × 0,28 × 0,7 μm (Blastocysts) and 0.23 × 0.23 × 1 μm (E6.5)

### Image analysis / cell polarity analysis

Nuclei in 3D confocal images were detected using either Nessys (Blin *et al*, 2019) for LaminB1 stained embryos or Farsight (Al-Kofahi et al., 2010) for propidium iodide stained pluripotent stem cells. The result of automated segmentations were then manually corrected using the Nessys editor tool.

Images were further analysed using PickCells (https://pickcellslab.frama.io/docs/). In order to detect centrosomes we used the difference of Gaussian method (Marr & Hildreth, 1980) implementation of ImgLib2 (Pietzsch *et al*, 2012). and manually corrected the result of the automated step using the PickCells 3D spot editor module. To create a polarity vector for each individual cell, centrosomes were assigned to their respective nuclei using the nucleus to centrosome assignment module in PickCells (documentation available at https://pickcellslab.frama.io/docs/use/features/segmentation/nc_assignments/).

The polarity angle was then defined as follows:

- For cell culture, the polarity angle was defined as the angle between the N/GC vector of the cell and the reference vector (0,0,−1) (vector pointing towards the bottom of the dish)
- For embryos, we first defined the embryo cavity manually in the 3D editor and then computed the vector normal to the surface of the cavity in PickCells. The polarity angle was defined as the angle between the cell’s polarity vector and the vector normal to the embryo cavity.

Angles were then exported to a csv file to be plotted either in R using the ggplot2 library (R Core Team; Wickham) (Fig. 1 beeswarm-boxplot) or with Rose.Net v0.10.0 (http://mypage.iu.edu/~tthomps/programs/home.htm).

The 3D representations were generated with PickCells (https://pickcellslab.frama.io/docs/)

## Supplemental Figures

**Supplemental Figure 1:**
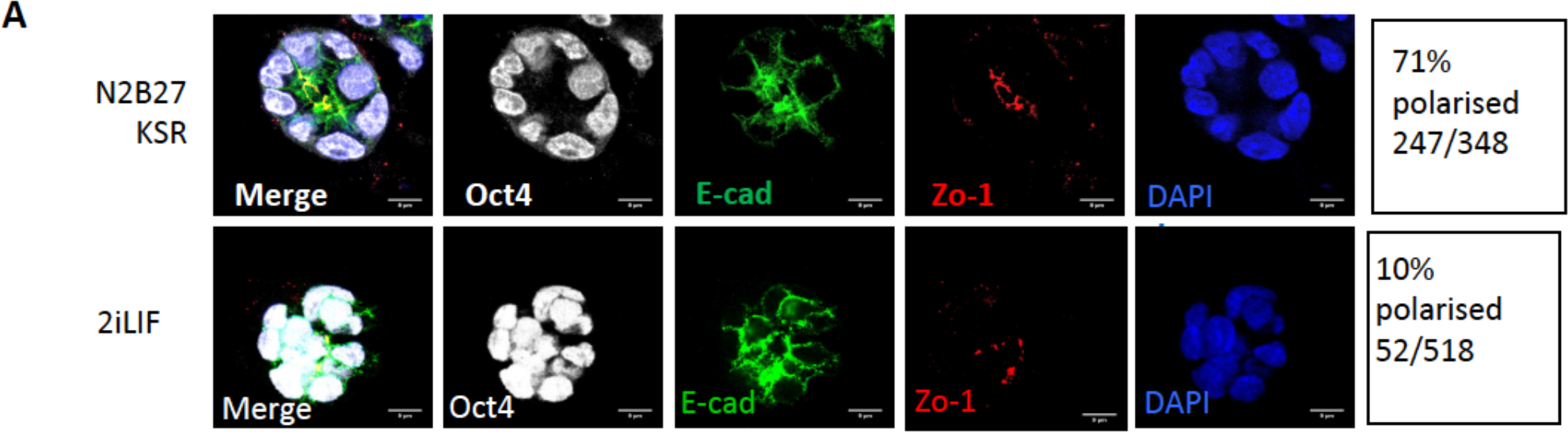
Naive pluripotent cells do not form stable polarised cysts. A: Cells were cultured for 48h in 600μg/ml matrigel in PBS at a density of 10,000 cells per well of a six-well plate in the presence of either N2B27+10%KSR for 48h (to enable differentiation) or 2iLIF(to block differentiation) then stained for the indicated markers.

**Supplemental Figure 2:**
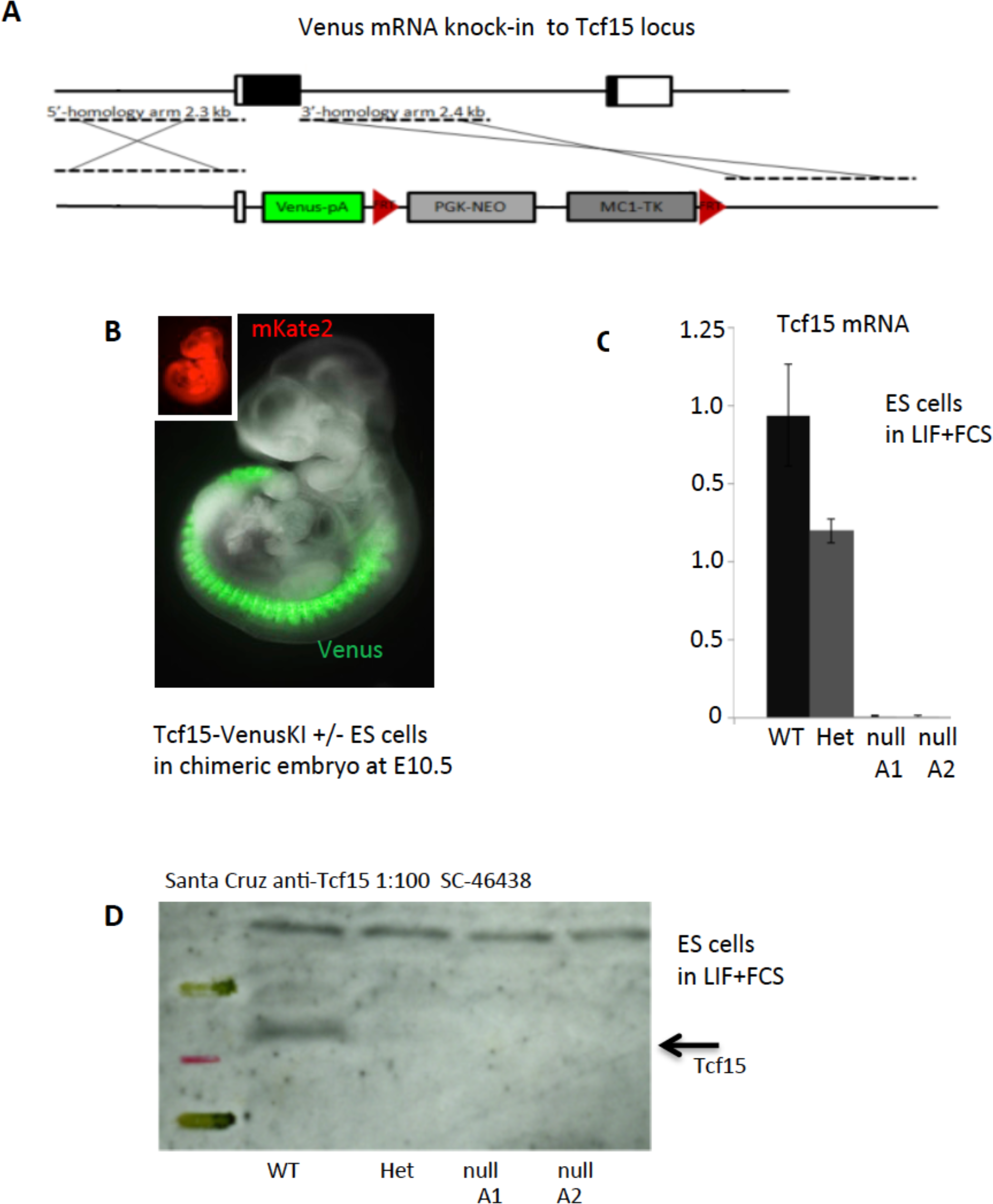
Generation and validation of Tcf15 VenusKI heterozygous and null cells. A: Targeting strategy for generating Tcf15VenusKI cells. The first coding exon of Tcf15 was replaced by the coding region of Venus in order to generate Tcf15-VenusKI +/− cell lines. A second round of targeting was performed in order to generate Tcf15-VenusKI −/− cell lines. B: Tcf15-VenusKI- +/− ES cells were constitutively fluorescently labelled by transfection with a plasmid encoding mKATE2 under the control of a CAG promoter then aggregated with wild type morulae, transferred into pseudopregnant female, recovered at E10.5 and assessed for chimerism. ES cells contributed extensively throughout the embryo (mKate fluorescence) but Venus expression was appeared to be restricted to the somites. C: qPCR analysis for Tcf15 mRNA in ES cells of stated genotypes grown in LIF+FCS. D: Western blot analysis of Tcf15 protein in ES cells of stated genotypes grown in LIF+FCS.

**Supplemental Figure 3.**
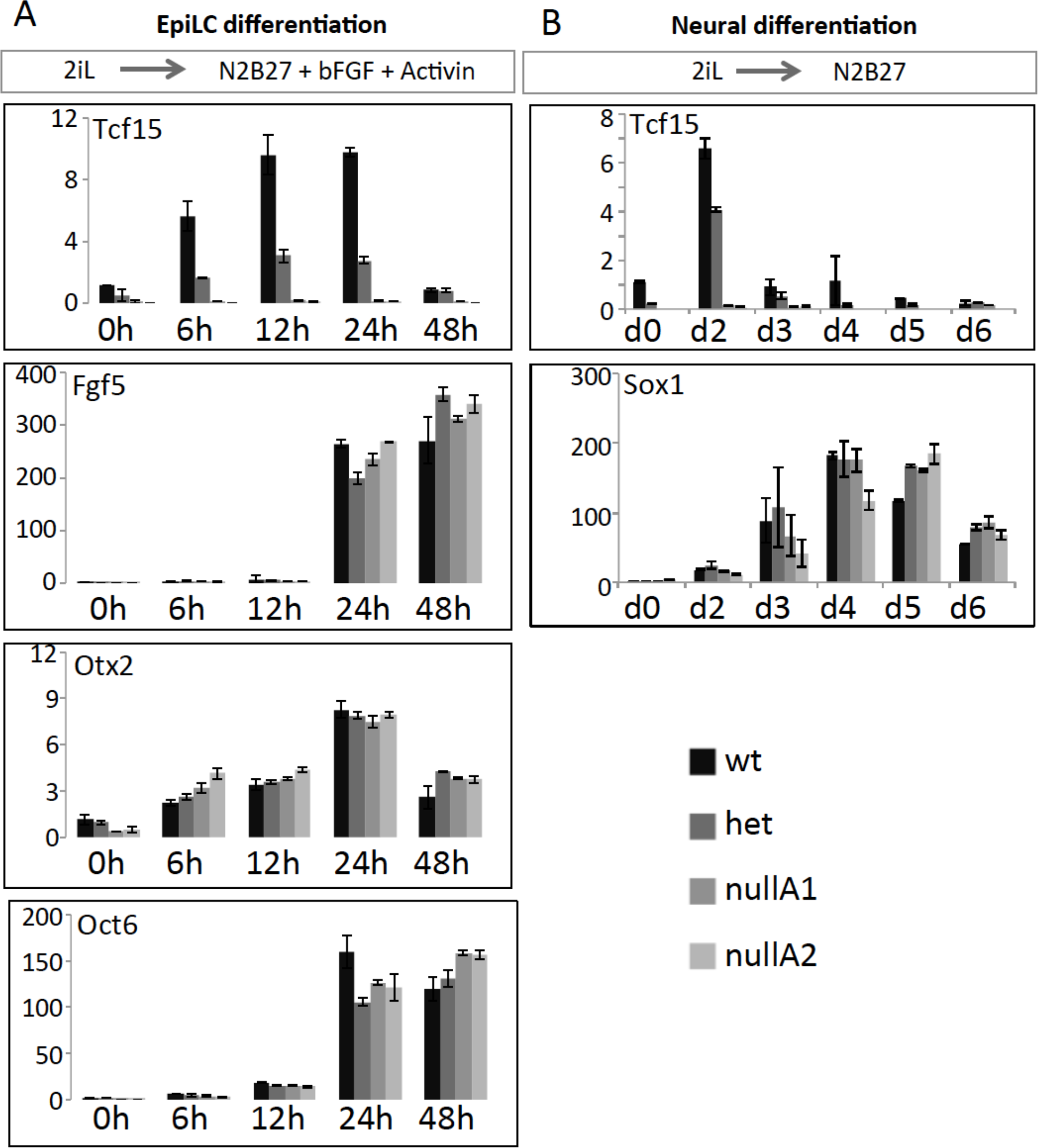
Tcf15 null cells do not have a differentiation phenotype in bulk culture. ES cells of stated genotypes subjected to EpiLC differentiation (A) or neural differentiation (B) then analysed by qPCR. Values are normalised to wild type cells at 0h.

